# Acquired epithelial WNT secretion drives niche independence of developing gastric cancer

**DOI:** 10.1101/2023.02.24.529854

**Authors:** Isaree Teriyapirom, Jihoon Kim, Heetak Lee, Sebastian R. Merker, Amanda Andersson-Rolf, Stephan R. Jahn, Anne-Marlen Ada, Sang-Min Kim, Joo Yeon Lim, Tim Schmäche, Nancy Wetterling, Saskia Stegert, Ji-Yeon Park, Jae-Ho Cheong, Hyunki Kim, Daniel E. Stange, Bon-Kyoung Koo

## Abstract

Recent studies have shed light on the signaling pathways required for gastric tissue maintenance and how aberrations in these key pathways lead to gastric cancer development. Although it has been shown that the WNT pathway is important for gastric epithelial homeostasis, the identity and source of the responsible canonical WNT ligands remain unknown. Furthermore, it is unclear how gastric cancer acquires WNT niche independence - an important early step in tumorigenesis. Using human and mouse gastric organoids and *in vivo* mouse models, we found that mesenchymal WNT2B and WNT7B maintain gastric epithelium in homeostasis. Next, mouse genetic studies and single-cell multi-omics analyses revealed that activation of MAPK signaling induces secretion of WNT7B in the epithelium itself. We further confirmed that in human gastric cancer, MAPK pathway activation through HER2 overexpression or copy number gains of *WNT2* confers WNT independence. Importantly, the epithelium-intrinsic WNT expression could be therapeutically inhibited. Taken together, our results reveal that normal gastric epithelial turnover relies on WNT ligands secreted by niche mesenchymal cells, while transformation involves acquisition of a WNT secretory phenotype in the epithelium - representing a potential target for therapeutic interventions.

## INTRODUCTION

In both humans and mice, the gastric corpus epithelium is continuously regenerated. The epithelium invaginates from the lumen to form glands, which are divided from top to bottom into four parts: pit, isthmus, neck and base. The murine gastric gland is maintained by two distinct stem cell populations, one situated in the isthmus and the other in the base ^1–5^. The isthmus stem cells (IsthSCs) are rapidly cycling and maintain the upper region of the gland ^4,6,7^. The cycling IsthSCs are marked by proliferation markers such as KI67 and STMN1, while the stem cells at the base (BSCs) are slow-cycling in homeostasis and marked by the WNT-signaling markers TNFRSF19 (TROY) and GPR49 (LGR5). These BSCs are normally gastric chief cells and function as reserve stem cells upon injury ^1,8,9^. The BSCs (or gastric chief cells) have also been identified as a source of gastric cancer ^9–13^.

The WNT signaling pathway plays a pivotal role in the maintenance of the gastrointestinal epithelium. For example, in small intestinal crypts, Paneth cells within the intestinal epithelium secrete Wnt3, driving self-renewal of neighboring stem cells and thus tissue maintenance ^14^. Mesenchymal cells under the intestinal crypt niche form another source of WNTs, secreting WNT2B as well as R-spondin 3 (RSPO3), a WNT signaling enhancer that binds to LGR4/5 expressed on intestinal stem cells (Gregorieff et al., 2005; de Lau et al., 2011; Farin, van Es and Clevers, 2012; Kang, Yousefi and Gruenheid, 2016).

Similar to the intestine, RSPO3 secreted by niche mesenchymal cells also plays an important role in regulating stem cell function in the stomach (Sigal et al., 2017; Fischer et al., 2022). WNT target genes such as AXIN2, LGR5, and TROY mark a stem cell population in the gastric gland base, and gastric stem cell-based organoid cultures require canonical WNT ligands such as WNT3A^1,8^. However, although WNT signaling is clearly essential for gastric stem cell function, the source(s) and the specific identity of the canonical WNT ligands present in the stomach are not well described.

A full understanding of the source and identity of WNT signals in the stomach is of major interest, as overactivation of WNT signaling, driving independence from a WNT niche environment, is often implicated in many epithelial cancers (Giles, van Es and Clevers, 2003; Anastas and Moon, 2012), of which gastric cancer is one of the most frequent and deadly ^19^. In the intestine, one of the best defined ligand-independent constitutive activation of WNT signaling caused by APC mutations, frees intestinal stem cells from the restricted niche space created by the WNT/RSPO gradient, and allows their continuous growth outside of the niche. This acquisition of niche independence is a key first step in intestinal and colonic tumorigenesis. In the gastric tissue, it has been shown that the loss of **<u>R</u>**NF43 and/or **<u>Z</u>**NRF3 (RZ), two negative regulators of the WNT receptors Frizzled, leads to independence from the WNT enhancer RSPO ^20^. RSPO independence can also be achieved by the combined loss of E-**<u>c</u>**adherin and T**<u>P</u>**53 (CP) ^21^, forming an alternative route. However, unlike APC mutations, RSPO independence *via* RZ or CP loss does not result in complete WNT independence, as WNT ligands are still needed to activate the pathway.

Here, we first used *in vivo* mouse models and gastric organoids to identify the specific WNT signaling molecules that drive maintenance of the gastric epithelium during homeostasis. Having identified WNT2B and WNT7B secreted by mesenchymal cells as the key WNT ligands in the stomach, we were then able to uncover two independent mechanisms that enable gastric epithelial stem cells to become fully WNT independent: MAPK-activation and genetic WNT gene amplification - both of which induce WNT secretion in the epithelium itself. Our results show that in contrast to colon cancer, WNT self-sufficiency in gastric cancer is established through different mechanisms that still require secretion and binding of WNT ligands – thus revealing a potential vulnerability of gastric cancer to WNT secretion blockers.

## RESULTS

### Gastric mesenchyme is the source of canonical WNT ligands in normal gastric epithelium

To identify the source of the canonical WNT ligands that maintain the gastric epithelium, we first separated the epithelium from the mesenchymal compartment and isolated RNA (Fig 1A). Successful separation of the compartments was confirmed by quantitative RT-PCR (qRT-PCR) for the mesenchymal marker BARX1 and the epithelial marker PGC (Fig 1B). Each compartment was then analyzed by qRT-PCR for expression of the *Wnt* gene family. The mesenchymal compartment expressed *Wnt2b, Wnt4, Wnt5a*, Wnt5b, *Wnt7b* and *Wnt9a*, while *Wnt* gene expression in the epithelial compartment was barely detectable (Fig 1C and Extended Data 1A). These data indicate that WNT is mainly secreted from gastric mesenchymal cells.

**Figure 1.**
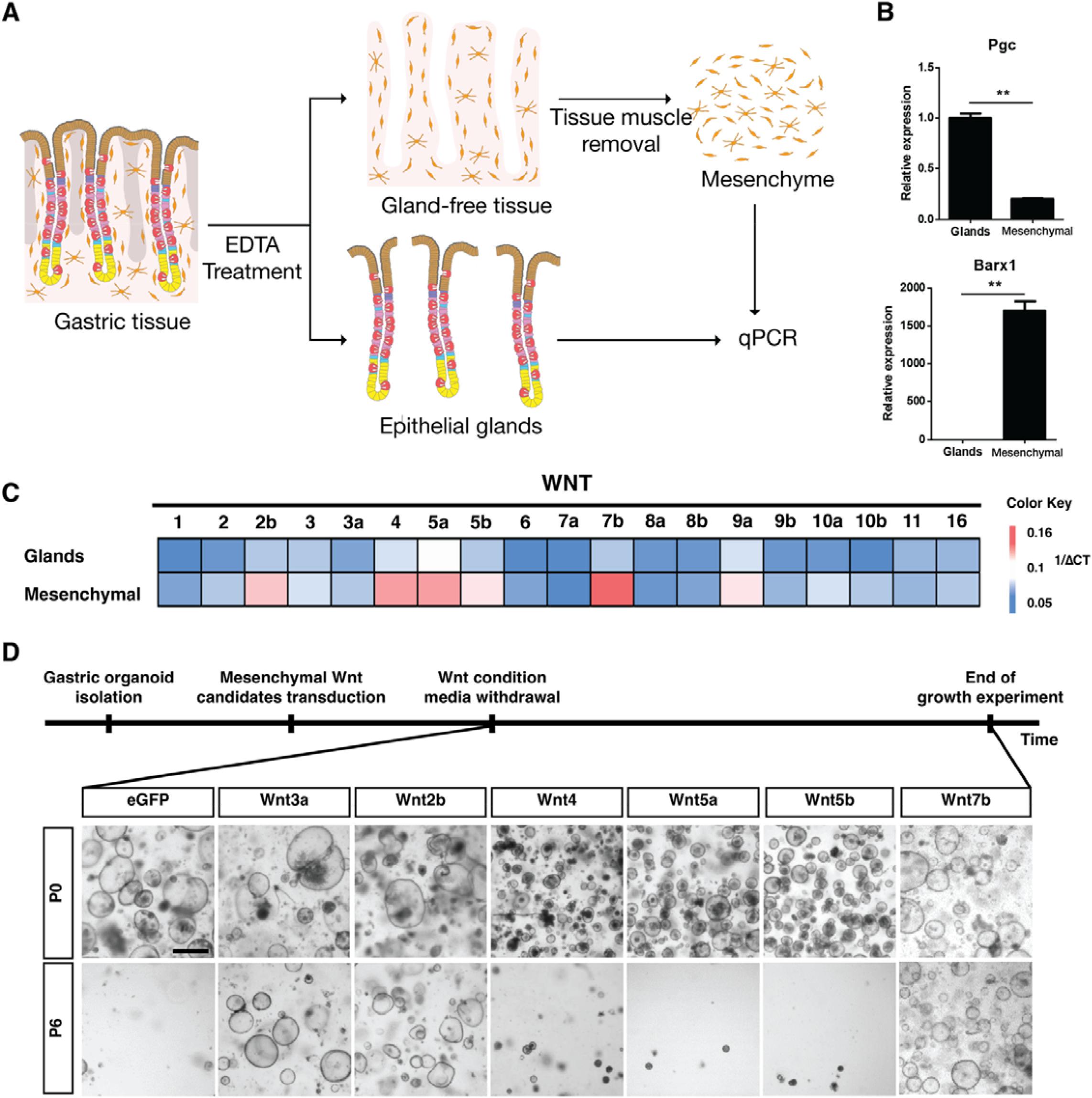
Gastric mesenchyme is the source of canonical WNT ligands in the normal gastric epithelium. (A) Experimental scheme for qRT-PCR analysis of gastric tissue following isolation of the epithelial glands from the mesenchymal component. (B) qRT-PCR of known markers of the epithelial gland (PGC) and mesenchymal (BARX1) compartment. (C) Heat map indicating expression of different WNT ligands in the epithelial gland and mesenchymal compartments. (D) Timeline schematic and representative organoid images of WNT retrieval assay. Gastric organoids were established from Rosa26-CreERT2 mice then transduced with retrovirus to overexpress a panel of candidate WNT genes identified in Fig 1C and Extended Data 1B,C. After tamoxifen treatment, WNT3A conditioned medium was removed, and organoid growth observed over six passages. P0, passage 0; P6, passage 6; scale bar, 1000 μm

We next utilized gastric organoids to functionally test the identified mesenchymal WNT ligands WNT2B, WNT4, WNT5A, and WNT7B for their ability to maintain turnover of epithelial stem cells *in vitro* (Fig 1D). We overexpressed candidate WNTs individually in gastric corpus organoids established from *Rosa26*-*Cre*^*ERT2*^ mice with a CRE-inducible retroviral overexpression system (Extended Data 1B and 1C)^22^. Overexpression of Wnt3A, a component of standard organoid medium, was used as a positive control, and eGFP as a negative control. After 4-OHT treatment to induce overexpression, infected organoids were cultured in WNT-deficient medium to test whether any of the overexpressed WNT candidates are able to replace the medium supplement WNT3A (Fig 1D). Overexpression of two mesenchymal WNT ligands, WNT2B and WNT7B, as well as the WNT3A control, were able to support the growth of gastric organoids in the absence of an external WNT source. The remaining ligands expressed in the mesenchyme, WNT4, WNT5A and WNT5B, were not able to support organoid growth, similar to the negative eGFP control. These data show that mesenchymal WNT2B and WNT7B are functionally relevant canonical WNT ligands for gastric stem cell maintenance.

### Acquisition of Kras mutation drives canonical WNT ligand production in the gastric epithelium

We and others have previously shown that loss-of-function mutations in RZ confer RSPO1 independence in the intestine and stomach, but this phenotype still depends on a paracrine WNT source ^20,21^. Since RNF43 mutations are also frequently found in gastric cancer ^2,23^, we decided to investigate the role of RZ loss of function in the stomach. To this end, we used *Anxa10-Cre*^*ERT2*^; *Rnf43*^*f/f*^; *Znrf3*^*f/f*^ (Ax10-RZ) mice (Extended Data 2A, 2B, 2C), which allow stomach-specific inducible deletion of RZ ^24^. One month after tamoxifen injection, we observed hyperplasia of the gastric glands, but with a normal gland architecture with a proliferating isthmus zone, as compared to wild-type control (CTRL) (Fig 2A). Since the MAPK pathway is also frequently activated in gastric cancer, we next included this in our model by adding a KRAS mutation (Kras^G12D^) in the background of the RZ loss-of-function (*Anxa10-Cre*^*ERT2*^: *Rnf43*^*f/f*^: *Znrf3*^*f/f*^: *Kras*^*lsl-G12D*^ (Ax10-RZK). One month after induction, we observed enhanced tumorigenic activity in the stomach epithelium compared to the Ax10-RZ mice, with extensive epithelial thickening and transformation (H&E) and widespread cellular proliferation (KI67) (Fig 2A). Despite the histological phenotype, it was still unclear whether the introduction of a KRAS mutation confers additional niche independence.

**Figure 2.**
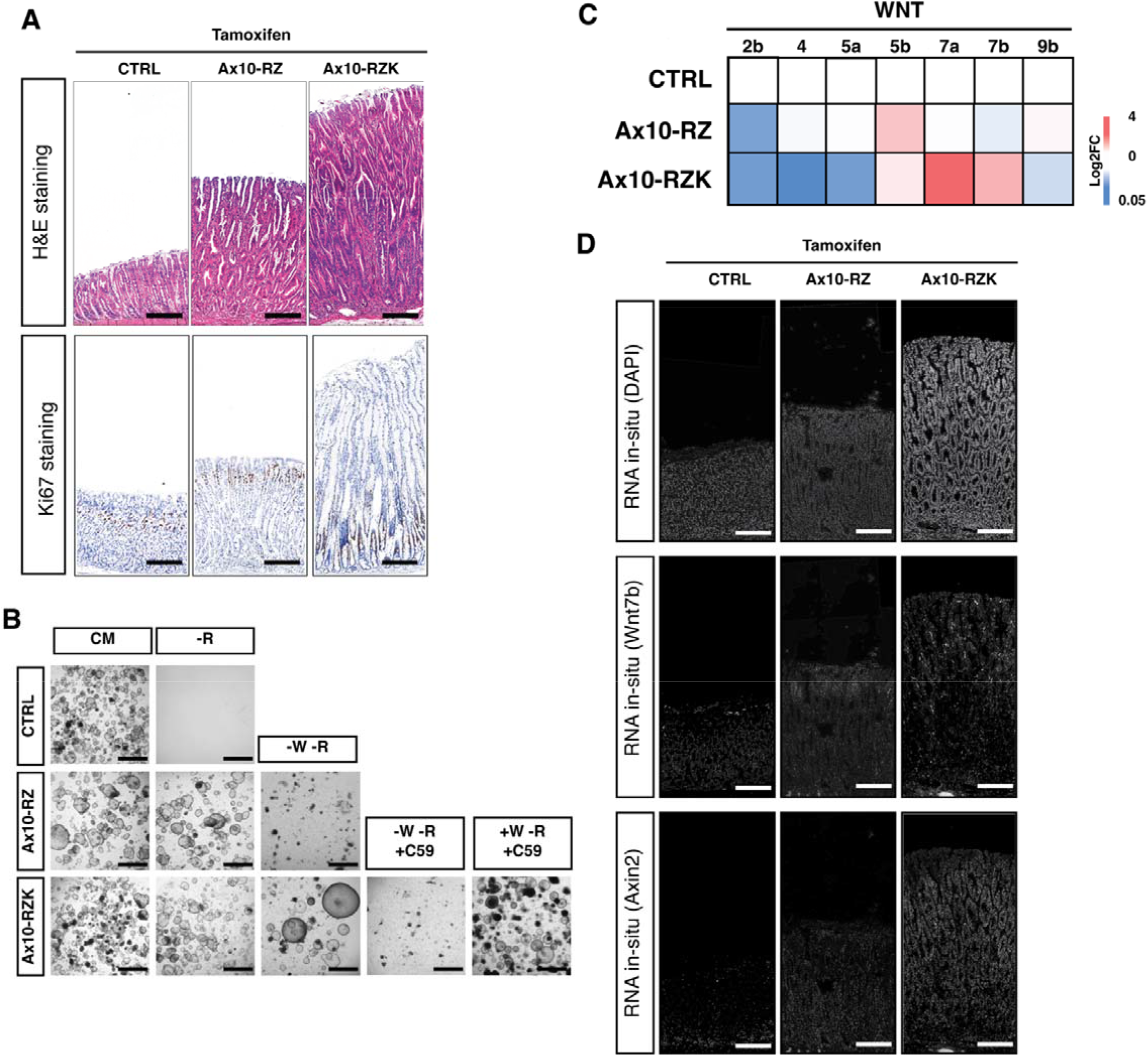
Acquired KRAS mutation drives canonical WNT secretion in the gastric epithelium. (A) Immunohistochemistry of corpus epithelium from control (CTRL), Anxa10-Cre^ERT2^: Rnf43^f/f^: Znrf3^f/f^ (Ax10-RZ) and Anxa10-Cre^ERT2^: Rnf43^f/f^: Znrf3^f/f^: l2l-Kras^G12D^ (Ax10-RZK) mice 1 month after tamoxifen induction. Shown are representative images of 2-4 mice used per genotype. Scale bars, 100 μm (B) Niche requirements of CTRL, Ax10-RZ and Ax10-RZK gastric organoids. Organoid growth was examined after 6 passages. Healthy organoids are cystic with a clear center. Shown are representative images of organoids isolated from 2-4 mice per genotype. CM, complete medium; -R, WENF; -W-R, ENF; -W-R+C59, ENF medium with C59 (10 mM); +W-R+C59, WENF with C59 (10 mM). Scale bars, 1000 μm (C) Heat map indicating expression of a panel of candidate WNT ligands (identified in Fig 1C) determined by qRT-PCR of isolated RNA from CTRL, Ax10-RZ and Ax10-RZK organoids. Expression is normalized to CTRL. n=3 (D) Representative multiplexed in situ hybridization images of CTRL, Ax10-RZ, and Ax10RZK mouse gastric tissue. Scale bars, 100 μm

To investigate niche independence phenotype, we next used gastric organoid cultures, where the culture medium acts as a proxy for the *in vivo* niche environment, providing essential growth factors, including EGF, NOGGIN, FGF10, WNT3A and RSPO1 ^1,8^. Depleting the culture of individual growth factors thus allows assessment of progressive niche independence of organoids derived from different stages of tumor progression. To determine whether the histological phenotypes of Ax10-RZ and Ax10-RZK correspond to niche independence, gastric corpus organoids from wild-type control, Ax10-RZ and Ax10-RZK mice were subjected to growth factor withdrawal experiments. As expected, while control organoids only grew in complete medium (CM), both Ax10-RZ and Ax10-RZK organoids showed sustained organoid growth in the absence of RSPO1, indicating independence from an external RSPO1 source ^20,21^ (Fig 2B). The Ax10-RZK organoids also showed strong EGF/FGF independence, confirming that MAPK signaling is constitutively active (Extended Data 2D). In contrast to Ax10-RZ organoids, Ax10-RZK organoids continued to grow even when WNT3A and RSPO1 were simultaneously withdrawn from the culture medium (Fig 2B). Collectively, these results indicate that oncogenic KRAS activation not only stimulates the MAPK pathway but also confers WNT independence.

As we observed a WNT independence phenotype in the Ax10-RZK organoids, we wanted to know whether this WNT3A independence was due to an acquired ability to self-secrete WNT ligands. To test this, we added C59, a Porcupine inhibitor that blocks WNT secretion, to culture medium depleted of WNT3A and RSPO. This resulted in growth arrest of Ax10-RZK organoids, which could be overcome by re-exposure to exogenous WNT3A (Fig 2B). These data indicate that the survival of Ax10-RZK gastric organoids in the absence of exogenous WNT3A was dependent on auto-secretion of canonical WNT ligand(s) from the epithelial cells of the gastric organoid. This confirms the existence of an unexpected link between MAPK activation upon oncogenic KRAS expression and WNT ligand secretion in the gastric epithelium.

To determine which specific WNT ligands are responsible for the auto-secretion phenotype, we compared expression levels of selected *Wnt* genes, previously identified as expressed in the mesenchymal compartment of the gastric tissue (Fig 1C) in Ax10-RZ and Ax10-RZK organoids using qRT-PCR. We observed a higher expression of *Wnt5b, Wnt7a* and *Wnt7b* in Ax10-RZK organoids (Fig 2C). While WNT5A/B are known ligands for non-canonical WNT signaling, WNT7B has shown ligand activity for the canonical WNT pathway in our gastric organoid overexpression experiments (Fig 1D). *In situ* hybridization also revealed significantly extended region of WNT7B and WNT downstream target gene AXIN2 expression throughout the gastric epithelium of Ax10-RZK compared to Ax10-RZ or control mice (Fig 2D). Thus, we conclude that WNT7A/B is the likely epithelial ligands for WNT signaling upon MAPK pathway activation.

### KRAS-mediated MAPK activation signals through the SPEM/Proliferative population to drive WNT7B production

To determine the mechanism through which activation of the MAPK pathway leads to autonomous secretion of WNT7B in the gastric epithelium, we performed single-cell multi omics analysis (scMultiomics incl. scRNA-seq and scATAC-seq) on Ax10-RZ and Ax10-RZK gastric organoids. In total, 10,809 high-quality cells from the two conditions were first visualized in a combined uniform manifold approximation and projection (UMAP) (Fig 3A). We identified 6 distinct clusters of epithelial cells by unsupervised clustering and annotated them based on the expression patterns of known marker genes for different gastric gland cell populations (Fig 3A, B). We detected not only the differentiated cell populations of pit (P) and neck (N) cells, but also several progenitor populations such as Lgr5+ (L), SPEM+ (S), and proliferating (Pr) cells. Of note, we also identified a novel population characterized by high levels of WNT7A/B expression, which we classified as Wnt7 (W).

**Figure 3.**
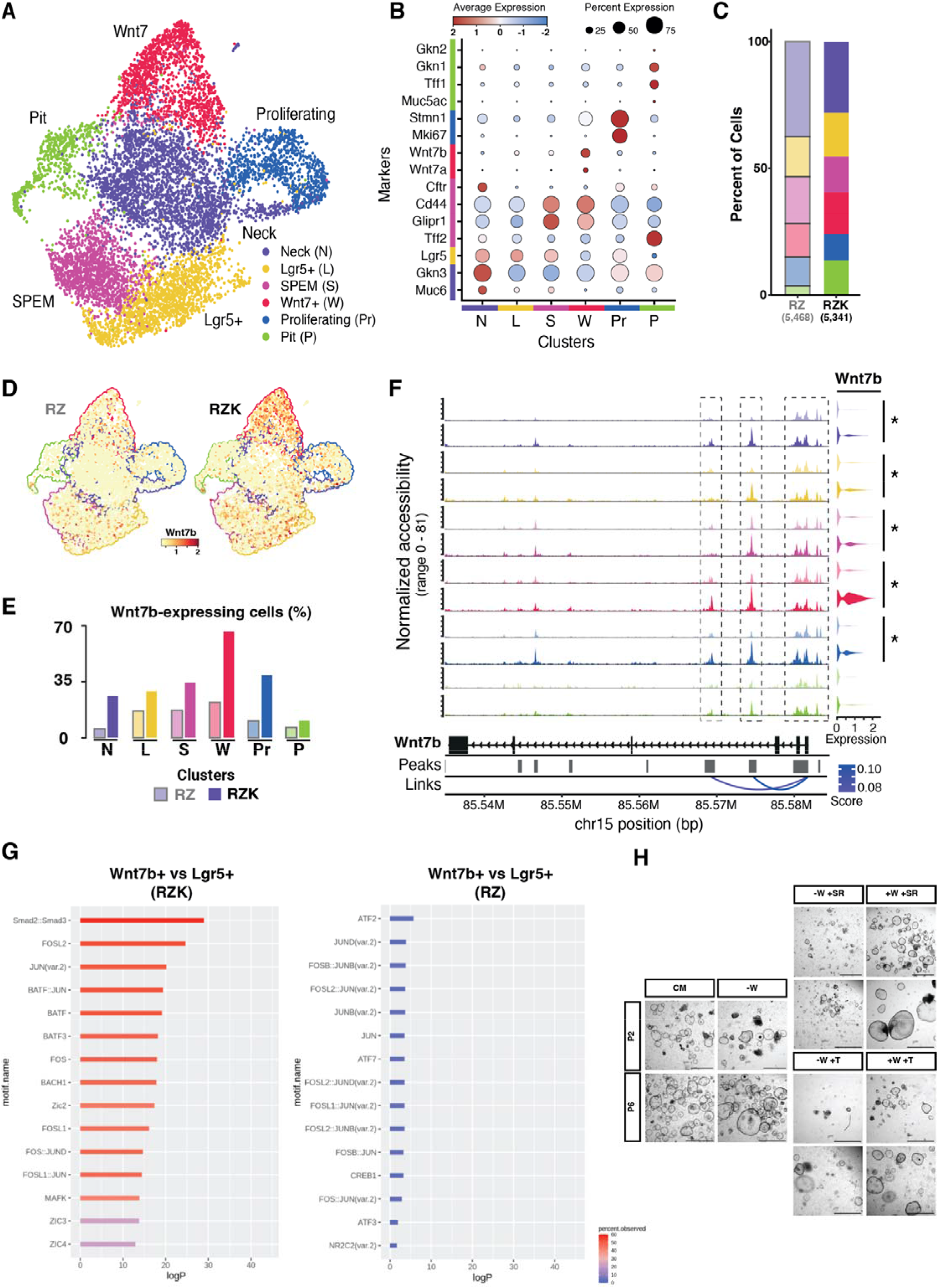
KRAS-mediated MAPK activation primes SPEM/LGR5+ population for differentiation to WNT7B producing cells. (A) Integrative uniform manifold approximation and projection (UMAP) plot showing clustering of epithelial cells from Ax10-RZ and Ax10-RZK organoids based on marker expression: neck, LGR5+, SPEM, WNT7+, Proliferating, and Pit cells. (B) Dot plots for expression of marker genes in each cell type. Average expression levels are indicated by color, and number of expressing cells by size of the dots, as indicated at the top. RZ, Ax10-RZ; RZK, Ax10-RZK; N, neck; L, LGR5+; S, SPEM; W, WNT7+, Pr, proliferating; P, pit. (C) Fraction of the different epithelial cell types out of the total cells analyzed in Ax10-RZ (lighter colors) and Ax10-RZK (darker colors) organoids. RZ, Ax10-RZ; RZK, Ax10-RZK. (D) UMAP plots showing WNT7B gene expression in Ax10-RZ and Ax10-RZK organoids. RZ, Ax10-RZ; RZK, Ax10-RZK. (E) Percentage of WNT7B-expressing cells in the different epithelial cell clusters of Ax10-RZ (lighter colors) and Ax10-RZK (darker colors) organoids. RZ, Ax10-RZ; RZK, Ax10-RZK; N, neck; L, LGR5+; S, SPEM; W, WNT7+, Pr, proliferating; P, pit. (F) ATAC-seq signal in different epithelial cell clusters of Ax10-RZ (lighter colors) and Ax10-RZK (darker colors) organoids in the WNT7B locus. Colors represent different cluster identities: purple, neck; yellow, LGR5+; pink, SPEM; red, WNT7+; blue, Proliferating; green, Pit, as shown in A. (G) Comparison of active transcription factors of the AP-1 transcription factor complex of Wnt7+ and Lgr5+ clusters from Ax10-RZ and Ax10-RZK. Motif enrichment analysis from differential accessible peak analysis using data from scMultiomics in the Wnt7+ as compared to Lgr5+ cluster in the Ax10-RZK (left panel) and Ax10-RZ (right panel). Enrichment is depicted in percentage observed on a logP scale. (H) AP-1 inhibitor treatment on CTRL, Ax10-RZ and Ax10-RZK gastric organoids. Organoid growth was examined after 6 passages with drug treatment. Healthy organoid growth is cystic with a clear center. Shown are representative images of organoids isolated from 2-4 mice per genotype. CM, complete medium; -W-R, ENF; -W-R+S, ENF medium with SR-11302 (50 μM); +W-R+S, WENF with SR-11302 (50 μM); -W-R+T, ENF medium with T-5224 (50 μM); +W-R+T, WENF with T-5224 (50 μM). Scale bars, 1000 μm.

When comparing Ax10-RZ and Ax10-RZK gastric organoids, the cell clusters and numbers remained relatively stable (Fig 3C). Nevertheless, in Ax10-RZK organoids, we observed increased expression of *Wnt7a/b*, mainly in the W cluster but at lower levels also in the S and Pr clusters (Fig 3D, E).

Next, we utilized the single-cell ATAC-seq profiles of the different clusters from our scMultiomics data set to compare DNA accessibility of the Wnt7b locus across the genome of Ax10-RZ (light colors) and Ax10-RZK (dark colors) organoids (Fig 3F). The Wnt7b locus shows eight distinct peak positions, indicative of transcription factor binding sites and open chromatin regions. Compared to Ax10-RZ, Ax10-RZK gastric organoids showed significantly higher peaks in the *Wnt7b* gene locus in all clusters, denoting more open chromatin (Fig 3F). Interestingly, the increased openness was already apparent in progenitor populations e.g. Lgr5+ (L), SPEM+ (S), and proliferating (Pr) cells, and was further enhanced in the W cluster where high-level of *Wnt7b* expression was observed (Fig 3F). These data suggest that KRAS activation already initiates opening of *Wnt7b* in progenitor cells, which then results in robust epithelial WNT production from the WNT-secreting cells (W cluster).

By analyzing differentially enriched transcription factors (TFs) of Ax10-RZ and Ax10-RZK gastric organoids between Lgr5+ and Wnt7 clusters, we identified that the well-known MAPK downstream TFs – FOS and JUN – as potential mediators of enhanced WNT7A/B expression and secretion in the Ax10-RZK organoids upon KRAS activation (Fig 3G). Upon heterodimerization, FOS and JUN form the AP-1 complex ^25^. We thus treated Ax10-RZK gastric organoids with the c-FOS/AP-1 inhibitors T-5224 or SR-11302 ^26–28^ and found that growth was significantly suppressed in the absence of WNT3A and RSPO1 (Fig 3H). Of note, treatment with SR-11302 caused a more severe growth inhibition than T-5224 treatment. Re-exposure to WNT3A rescued the growth of Ax10-RZK organoids in the presence of the c-FOS/AP-1 inhibitors. Taken together, these results show that mutation of KRAS facilitates the opening of *Wnt7a/b* chromatin through the KRAS-MAPK-AP-1 axis.

### WNT independence can be acquired by divergent pathways

Following our finding that KRAS-mediated MAPK activation coupled with RZ loss-of-function leads to a WNT secretion phenotype in the gastric epithelium, we asked whether this phenotype depends specifically on the combination of these three mutations (RZK). It has been shown that the gastric epithelium can obtain RSPO1 independence not only by losing RZ function but also by combined loss of E-**c**adherin (*Cdh1*) and T**P**53 (*Tp53*) (CP) ^21^. To model this alternative path to RSPO1 niche independence, we generated two further conditional knockout mouse lines based on Anxa10-Cre^ERT2^, using the alleles *Cdh1*^*f/f;*^ *Tp53*^*R172H*^ (Ax10-CP) and in addition Kras^l2l-G12D^ (Ax10-CPK) (Extended Data 3A and 3B). Following tamoxifen induction, we observed gastric epithelial hyperplasia and cellular proliferation in Ax10-CP mice, which was further enhanced in the Ax10-CPK model (Fig 4A). Furthermore, upon growth factor withdrawal gastric organoids established from wild-type, Ax10-CP and Ax10-CPK mice demonstrated similar phenotypes as observed above with the Ax10-RZ and Ax10-RZK mice: both survived over multiple passages in medium without RSPO1 but only Ax10-CPK organoids maintained continuous growth when both WNT3A and RSPO1 were withdrawn (Fig 4B). Thus, the CPK organoids reproduced the same WNT self-sufficient phenotype as seen in RZK organoids (Fig 2B). This was further validated by treating Ax10-CPK organoids with the Porcupine inhibitor C59 (Fig 4B). Ax10-CPK organoids could not be maintained in the presence of C59 but were rescued by re-exposure to WNT3A. In order to identify the secreted WNT ligands that maintain growth in the absence of WNT3A, we used qRT-PCR to compare the expression of candidate WNT genes in Ax10-CP and Ax10-CPK organoids (Fig 4C). Interestingly, only *Wnt7a/b* were highly upregulated in Ax10-CPK organoids, implicating a similar mechanism of WNT7A/B auto-secretion by gastric epithelial cells as in the Anx10-RZK model (Fig 2C). Taken together, these results suggest that WNT independence in the gastric epithelium through the KRAS-induced self-secretion of WNT ligands is not limited to cells with RZ loss of function but can also be achieved by CP mutations.

**Figure 4.**
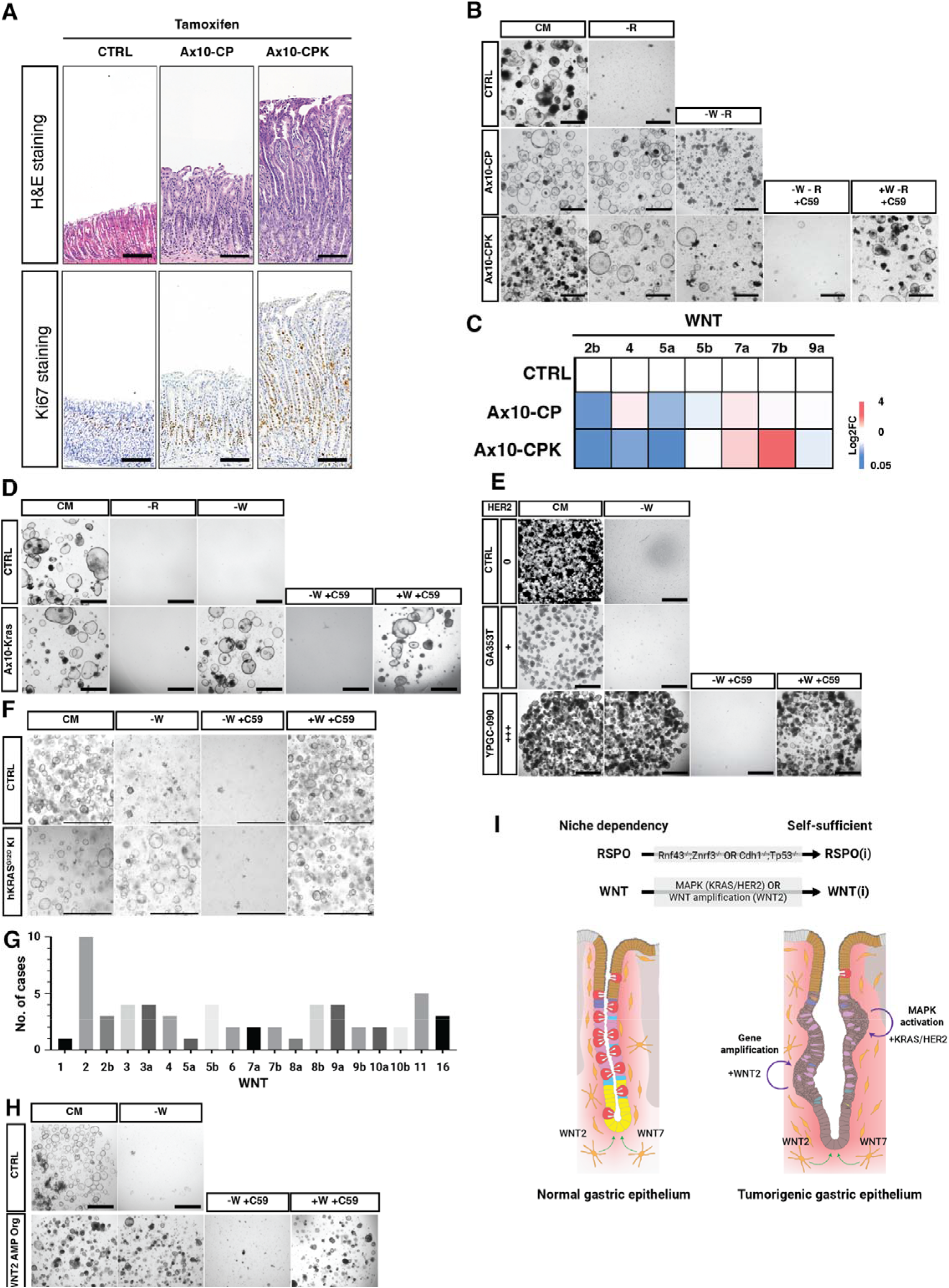
Alternative pathways to obtaining WNT independence. (A) Immunohistochemistry of corpus epithelium from control (CTRL), Anxa10-Cre^ERT2^: Cdh1^f/f^: Tp53^f/f^ (Ax10-CP) and Anxa10-Cre^ERT2^: Cdh1^f/f^: Tp53^f/f^: l2l-Kras^G12D^ (Ax10-CPK) mice 1 month after tamoxifen induction. Shown are representative images of 2-4 mice used per genotype. Scale bars, 100 μm (B) Niche requirements of CTRL, Ax10-CP and Ax10-CPK gastric organoids. Organoid growth was examined after 6 passages. Healthy organoid growth is cystic with a clear center. Shown are representative images of organoids isolated from 2-4 mice per genotype. CM, complete medium; -R, WENF; -W-R, ENF; -W-R+C59, ENF medium with C59 (10 mM); +W-R+C59, WENF with C59 (10 mM). Scale bars, 1000 μm. (C) Heat map indicating expression of a panel of candidate WNT ligands (identified in Fig 1C) determined by qRT-PCR of isolated RNA from CTRL, Ax10-CP and Ax10-CPK organoids. Expression is normalized to CTRL. n=3 (D) Niche requirements of CTRL and Anxa10-Cre^ERT2^: lsl-Kras^G12D^ (Ax10-Kras) gastric organoids. Organoid growth was examined after 6 passages. Healthy organoid growth is cystic with a clear center. Shown are representative images of organoids isolated from 2-4 mice per genotype. CM, complete medium; -R, WENF; -W, ENFR; -W +C59, ENFR medium with C59 (10 mM); +W+C59, WENFR with C59 (10 mM). Scale bars, 1000 μm. (E) Niche requirements of human gastric cancer patient-derived organoid lines with HER2 activation. HER2 levels are classified as negative (0), low-grade (+) and positive (+++) based on guidelines published by Bartley et al., 2016. Healthy organoid growth is cystic with a clear center or growing in grape-like structures. CM, complete medium; -W, ENFR; -W+C59, ENFR medium with C59 (10 mM); +W+C59, WENFR with C59 (10 mM). Scale bars, 1000 μm. (F) Niche requirements for human gastric organoid lines (WT) and organoids targeted with human Kras^G12D^ knock-in (hKRAS^G12D^KI). Healthy organoid growth is cystic with a clear center. CM, complete medium; -R, WENF; -W-R, ENF; -W-R+C59, ENF medium with C59 (10 mM); +W-R+C59, WENF with C59 (10 mM). Scale bars, 1000 μm. (G) TCGA analysis of WNT gene amplification occurrences in gastric cancer samples. (H) Niche requirements of human gastric organoid lines (WT) and human gastric cancer patient-derived organoid lines with WNT2 gene amplification (hWNT2-Amp). Healthy organoid growth is cystic with a clear center or growing in grape-like structures. CM, complete medium; -R, WENF; -W-R, ENF; -W-R+C59, ENF medium with C59 (10 mM); +W-R+C59, WENF with C59 (10 mM). Scale bars, 1000 μm. (I) Schematic model showing WNT requirements for normal gastric epithelial and acquired mutations driving independence from external niche sources in tumorigenesis. Inset depicts a model mechanism where SPEM/LGR5+ cells (pink, yellow cells, respectively) receive MAPK signaling, eventually differentiating into WNT7+ cells (red cells).

Since the presence of RZ or CP mutations alone already conferred RSPO1 independence in organoid cultures derived from Ax10-RZ and Ax10-CP mice, (Fig 2B and Fig 4B), this raised the question of whether WNT independence can only be obtained subsequent to RSPO1 independence or also directly by the acquisition of MAPK activation. To investigate this question, we generated gastric organoids from *Anxa10*-Cre^ERT2^; *Kras*^*lsl-G12D*^ (Ax10-Kras) mice and compared their growth to organoids from wild-type mice (Fig 4D). While wild-type organoids died when any growth factor (i.e. either RSPO1 or WNT3A) was removed, Ax10-Kras organoids were able to grow without WNT3A but died without RSPO1 (Fig 4D). Treatment with C59 inhibited the growth of Ax10-Kras organoids in WNT3A-free medium, which was rescued by re-introduction of WNT3A to the medium (Fig 4D). WNT independence is therefore not an event that is secondary to R-SPONDIN independence, but both occur independently from each other, forming divergent routes to niche independence in gastric cancer development.

### MAPK activity and WNT2 copy number gain in gastric cancer patient-derived organoids correlate to WNT independence

Next, we set out to investigate whether the identified mechanisms of WNT independence in mice are also relevant to human gastric cancer development. Gene amplification and subsequent overexpression of HER2, a receptor tyrosine-protein kinase acting upstream of the KRAS-MAPK pathway, are commonly found in gastric cancers. Therefore, we analyzed gastric cancer patient-derived organoids (GC-PDOs) from primary cancers with different levels of HER2-positivity according to standard immunohistochemistry testing (Fig 4E and Extended Data 3C). The GC-PDO line GA353T, expressing only mildly elevated levels of HER2, was unable to grow in the absence of WNT3A in the culture medium (Fig 4E). In contrast, GC-PDO lines with high HER2 expression levels (YPGC-090, GA372T, YPGC-021, -075, and -105) could be maintained without WNT3A (Fig 4E and Extended Data 3D). Inhibition of WNT secretion by C59 prevented growth, which could be rescued by re-addition of WNT3A to the medium (Fig 4E and Extended Data 3D). This suggests that HER2-mediated MAPK activity also confers WNT independence through self-secretory mechanism in human gastric cancer.

As GC-PDOs show extensive molecular alterations in addition to HER2 overexpression, which could potentially also affect their dependence on WNT, we utilized CRISPR/Cas9 gene editing to knock-in the KRAS^G12D^ mutation in a normal human gastric organoid line (hKRAS^G12D^-KI). Unlike the parent organoid line, hKRAS^G12D^-KI organoids were able to grow in the absence of WNT3A (Fig 4F). The organoid growth could be inhibited by the addition of C59 and rescued upon re-introduction of WNT3A to the medium. Thus, also in human gastric epithelial cells, WNT independence could be achieved by activation of MAPK signaling, phenocopying the observations made in our mouse models.

In order to identify further mechanisms through which gastric cancers can acquire WNT independence, we searched the TCGA dataset for WNT gene alterations in human gastric cancers. WNT gene copy number alterations were observed in all 19 WNT genes, with WNT2 being the most frequently altered (Fig 4G). WNT2B, a WNT2 paralogue, was also the canonical WNT ligand we found to be expressed in the mouse gastric mesenchyme and to be functionally able to maintain gastric epithelial proliferation (Fig 1C and 1D). From our cancer organoids, we identified a GC-PDO line with a WNT2 copy number gain (WNT2 AMP Org). This line was able to maintain growth in medium without WNT3A (Fig 4H), which was once again prevented by addition of C59 and could be rescued by re-addition of WNT3A (Fig 4H). This observation suggests an alternative route to MAPK pathway activation to achieve intrinsic WNT independence: direct upregulation of WNT ligand expression by copy number gains.

Taken together, we show that during homeostasis, turnover of the gastric epithelium is maintained by WNT2B and WNT7B secreted by the mesenchyme (Fig 4I). In gastric tumorigenesis, the acquisition of MAPK pathway activation by either KRAS mutation or HER2 overexpression, functioning through AP-1, induces expression and secretion of WNT7A/B in the gastric epithelium, thereby conferring WNT self-sufficiency to the epithelium itself. Alternatively, intrinsic Wnt gene overexpression, e.g. by WNT2 gene copy number gain, is another frequent event in gastric cancer, and could be functionally validated to result in WNT independence. Thus, developing gastric cancer can take divergent routes to WNT independence as an early but decisive step in tumor progression.

## DISCUSSION

While it has been shown that WNT activity is important for the function of gastric stem cells, the source of WNT and the identity of the functionally relevant WNT ligands for gastric tissue maintenance has not yet been adequately resolved. Using gastric epithelial organoids and WNT ligand profiling, here we identified the mesenchymal niche as the only compartment which secretes WNT ligands during gastric epithelial homeostasis. Functionally important WNT ligands for gastric tissue maintenance are WNT2B and WNT7B. Furthermore, we showed that the acquisition of MAPK pathway activation (by KRAS mutation or HER2 overexpression) as well as WNT2 gene copy number gains in human gastric cancer constitute an important step in the process of tumorigenesis by freeing the gastric epithelial cells from their dependence on a niche WNT source.

Tumorigenesis is often described as a step-by-step process, where one mutation or a combination of mutations leads to an initial transformation process that allows the cells to escape their dependence on external growth factors by acquiring mutations that activate the downstream signaling pathways. Colon cancer is a prime example of this dogma. The initial hit in form of an APC loss-of-function mutation initiates tumorigenesis in the colonic tissue by enabling the epithelium to be self-sufficient, losing its reliance on the external WNT and R-SPONDIN ligand source ^29,30^. This initiates benign adenoma formation, which can last many years until the transformed cells acquire a second oncogenic hit in the form of a KRAS^G12D^ mutation. This renders tumor cells independent from another external ligand, EGF, resulting in further progression to an intermediate adenoma stage. With subsequent mutations, the transformed adenoma cells then turn into full-blown cancer.

Although the two tissues are closely related, tumorigenesis in the gastric epithelium shows more divergence. Independence from niche-derived R-SPONDIN can be achieved in two different ways: either *via* RNF43 and ZNRF3 loss-of-function or E-cadherin and TP53 loss-of-function ^20,21^. WNT independence is acquired through activation of the MAPK pathway by KRAS activation and HER2 receptor upregulation or alternatively, direct upregulation of a WNT family gene e.g., by copy number gains. As we demonstrate here, WNT and R-SPONDIN independence in the gastric tissue also occur independently of each other, not sequentially or simultaneously as in the colon through APC loss-of-function. The crosstalk we identify between the two major signaling pathways MAPK and WNT in gastric cancer tumorigenesis potentially contributes to tumor cell plasticity, resulting in treatment failure when only one of the two pathways is therapeutically targeted. Our work suggests that a combination of WNT secretion inhibitors with targeted MAPK therapeutics could constitute a novel treatment approach for gastric cancer patients.

## MATERIALS AND METHODS

### Mice

All mouse lines were maintained in a controlled facility under the standard light/dark cycle, temperature, and humidity. All mouse experiments adhered to the guidelines of the Austrian Animal Care and Use Committee. The Villin-CreER^T2^: Rnf43^f/f^: Znrf3^f/f^ mice were generated previously ^31^. The Anxa10-CreER^T2^ line ^24^ used for the gastric epithelium-specific conditional genetic mutation Kras^lsl-G12D^ (MGI ID: 008179) was obtained from The Jackson Laboratory and crossed with Cdh1^f/f^: Tp53^R172H^ mice provided by Daniel Stange to generate Ax10-CP and Ax10-CPK lines.

### Animal treatments

To induce Ax10-CreER^T2^ activation, age-matched mutant mice together with wild-type negative control C57BL/6J mice were injected intraperitoneally with tamoxifen (Sigma, T5648) in corn oil at a concentration of 2 mg per 20g body weight at age 8 – 12 weeks. Both male and female mice were used for the experiments. One month after tamoxifen injection, mice were sacrificed by CO_2_ inhalation, and the stomach isolated for further histology and gland isolation.

### Murine stomach preparation for histology

Isolated stomach tissue was washed with cold PBS and the greater curvature cut longitudinally from the intestine to the esophagus. The sample was spread and secured with needles on a piece of cardboard before fixation in freshly prepared 4% PFA at 4 °C overnight (18 h) or in 10% NBF (neutral buffered formalin) overnight at room temperature while shaking. After fixation, the stomach tissue was washed with 1xPBS three times for 30 minutes at 4 °C.

### Paraffin embedding and Immunohistochemistry staining

For paraffin embedding, samples were dehydrated with ethanol (70, 80, and 100% for 80 minutes each) then with xylene and infiltrated in paraffin for three rounds of 100 minutes each. Samples were then embedded in a paraffin block and sectioned at 2 mm thickness for all histological analyses.

In preparation for immunohistochemistry, sections were first re-hydrated and the antigens were retrieved with sodium citrate (pH 6.0) following the VBC Histology Facility protocols.

For immunohistochemistry staining, all samples were incubated in blocking solution consisting of 3% H_2_O_2_ at room temperature for 10 min, followed by incubation in blocking solution consisting of 2% BSA, 5% goat serum, 0.3% Triton-X100 in PBS at room temperature for 1 hr. Recombinant anti-Ki67 primary antibody (1:200; Abcam; ab16667) was applied to each section and the peroxidase-conjugated 2-step enhancer-polymer system (DCS, SuperVision 2 HRP Single Species) was used for detection. Hematoxylin and eosin staining was carried out without heat using the Epredia Gemini AS Automated Slide Stainer.

RNA in situ hybridization to detect *Wnt7b* and *Axin2* was done using the RNAScope Multiplex Fluorescent Detection Kit v2 according to the manufacturer’s protocol (ACDBio323110). In brief, paraffin-embedded samples were sectioned to 4 μm thickness one week prior to staining. On the staining day, slides were manually pretreated to remove paraffin, retrieve target sequences and stored overnight. Staining was done according to the manufacturer’s protocol with probes for Wnt7b (Advanced Cell Diagnostics, Cat No. 401131) and Axin2 (Advanced Cell Diagnostics, Cat No. 400331-C3). Stained slides were imaged using the Pannoramic FLASK 250 III scanner (3DHISTECH) and images were processed using the CaseViewer software.

### Establishment and culture of mouse corpus epithelial gastric organoids

The mouse stomach glands were isolated as described previously^1,8^. Briefly, freshly isolated stomach was washed with cold PBS and the corpus region was separated. The corpus was cut into small pieces and incubated with Gentle Cell Dissociation Reagent (STEM Cell Technology) for 10 min. Isolated glands, were seeded in Matrigel (Corning) at a density of 100 – 150 glands per well and cultured in basal medium supplemented with growth factor cocktail. Basal medium consisted of advanced Dulbecco’s modified Eagle medium (DMEM)/F12, 1% penicillin/streptomycin, 10 mM HEPES (Gibco), 1% GlutaMAX (Gibco), 1x B27 (Life Technologies), 1.25 mM N-acetylcysteine (Sigma-Aldrich). Growth factors used for mouse gastric corpus organoid culture were 50 ng/ml mEGF (Peprotech), 100 ng/ml mNoggin (Peprotech), 10% R-spondin1 conditioned medium, 50% Wnt3A conditioned medium, 100 ng/ml hFGF10 (Peprotech), and 10 nM hGastrin (Sigma-Aldrich).

### Gastric glands and mesenchymal tissue separation

For identification of WNT ligand source, the stomach was harvested and epithelial glands were isolated using an EDTA treatment step. The remaining tissue was transferred to a Petri dish. Using two fine forceps, the white mesenchymal tissue was separated from the underlying muscle layers.

### WNT retrieval experiment with retroviral WNT overexpression in mouse gastric organoids

#### Retrovirus production

The retroviral infection system was used as previously described ^22^. Virus-producing PlatinumE cells (kind gift of Hans Clevers, Hubrecht Institute, Netherlands) were used to package and produce virus. Cells were thawed and washed two times in DMEM/F12 with 10% heat-inactivated FCS and 1% Pen/Strep (referred to as ++ medium) and centrifuged at 500 x g for 5 min between each washing step. The pellet was resuspended in 5 ml of ++ medium and plated in a 25cm^2^ cell culture flask (Corning). For selection purposes, puromycin (1 μg/μl) and blasticidin (10 μg/μl) were added, and cells incubated at 37 °C with 95% humidity and 5% CO^2^. Medium was replaced every 3 days and fresh antibiotics added. For passaging, cells were washed twice with PBS and detached by incubating in trypsin for 5 min at 37 °C. The trypsinization process was stopped by adding ++ medium. After centrifugation at 500 x g for 5 min, cells were either seeded for virus production or transferred into a bigger flask. For virus production, 0.8 × 10^7^ cells were seeded in 15 cm diameter Petri dishes and kept in 25 ml of ++medium without antibiotics. Cells were transfected using 30 μg of pMSCV-loxP-dsRed-loxP-eGFP-Puro-WPRE plasmid (Addgene, Plasmid #32702) or pMSCV-loxP-dsRed-loxP-Cited2-3HA-Puro-WPRE plasmid (Addgene, Plasmid #32703). DNA was mixed with Lipofectamin2000 (ThermoFisher) to a total volume of 250 μl and incubated for 30 min at room temperature. After incubation, 250 μl Opti-MEM (ThermoFisher) was added to the mix. This transfection mix was then carefully added to the Petri dishes containing the PlatinumE cells. One day post-transfection, PlatinumE cells were checked for successful transfection through detection of red fluorescence. Two days later, the virus-containing supernatant was collected, filtered through a 0.45 μm filter and centrifuged overnight at 8000 x g at 4 °C. The next morning, the supernatant was completely discarded, and the pellet resuspended in 20 μl infection medium. Infection medium consisted of mouse gastric organoid culture medium without pen/strep, and with primocin and 1:1000 polybrene (Sigma Aldrich) added. If the virus medium was not directly used, it was stored at -80 °C.

#### Retroviral infection of murine gastric organoids

Murine gastric organoids from the corpus region of Rosa26-Cre^ERT2^ mice were used for infection. 8 wells of a 48-well plate were pooled in a 15 ml tube and mechanically dissociated as described above for normal passaging. Cells were centrifuged at 300 x g and 4 °C for 5 min and the pellet resuspended in 500 μl Cell Recovery Solution (Corning) and kept for 10 min on ice. Cells were washed with 10 ml PBS and centrifuged for 5 min at 500 x g and 4 °C. The pellet was resuspended in 2 ml TrypLE (Gibco) and incubated for 2 min at 37 °C. The reaction was stopped with 2 ml infection medium and distributed in 3 × 15 ml tubes. The suspensions were then centrifuged at 500 x g and 4 °C for 5 min. In the meantime, the virus pellets (Cited2 and eGFP) were resuspended in 250 μl infection medium. Pellets were then resuspended in the corresponding virus solution or pure infection medium as control. Cell suspensions were plated on a 48-well plate, sealed with parafilm, and spinoculated at 32 °C for 60 min at 600 x g. After spinoculation, parafilm was removed and the plate incubated for 6 h at 37 °C, 95% humidity and 5% CO_2_. The content of each well was then transferred to a 15 ml tube, 1750 μl infection medium added, and the tube centrifuged at 300 x g and 4 °C for 5 min. Supernatant was discarded, the pellet resuspended in 20 μl Matrigel and plated in one well. After 15 min of incubation, 250 μl normal murine gastric organoid medium was added. 3 days post-infection, organoids were checked for red fluorescence under the microscope. Upon detection of red signals, puromycin selection was started. Puromycin was added at a concentration of 2 μg/ml and the selection was performed until the uninfected control organoids were dead. Once the selection was completed, organoids were induced using a 1:20,000 dilution of 5 μM hydroxy-tamoxifen (4-OHT; Sigma Aldrich) in mouse gastric organoid medium. Tamoxifen induction was performed for three days and the medium and 4-OHT changed daily.

### RNA purification and qRT-PCR

To confirm expression of individual *Wnt* genes in mouse and human gastric organoids, we extracted total RNA using the RNeasy Mini RNA Extraction Kit (Qiagen) according to the manufacturer’s instructions. Total RNA samples were analyzed by qRT-PCR using the primer sets against the *Wnt* genes listed in Supplementary Table 1.

### Establishment of patient-derived gastric cancer organoids

Human gastric cancer patient-derived organoids were established from tissues obtained from surgical procedure with informed consent and ethics permission from Yonsei University College of Medicine, Seoul, South Korea and maintained as described previously ^32^ and maintained in the following culture medium: AdDMEM/F12 with 1% penicillin/streptomycin, 10 mM HEPES (Gibco), GlutaMAX (Gibco), 50% Wnt3A conditioned medium, 10% R-SPONDIN conditioned medium, 100 ng/ml mNoggin (Peprotech), 1x B27 (Gibco), 10 mM nicotinamide (Sigma-Aldrich), 1.25 mM N-acetyl-L-cysteine (Sigma-Aldrich); 50 ng/ml hFGF10 (Preprotech), 50 ng/ml mEGF (Peprotech), 10 nM hGastrin (Sigma-Aldrich), 2 μM A83-01 (Tocris Bioscience), 0.5 mM PGE2 (Tocris).

#### HER2 3+ PDO

HER2-overexpressing organoids were established from patient-derived surgical specimens and ascites with informed consent and ethics permission from Yonsei University College of Medicine, Seoul, South Korea using a previously described protocol ^21^. HER2 activity was determined by the Pathology Department of the Yonsei University following an established guideline ^33^. The expression of HER2 was confirmed using immunohistochemistry in organoids and parent cancer tissues.

#### WNT2 amplification organoids

WNT2 gene amplification organoids were established according to the methods published in ^34^. Briefly, human gastric cancer and healthy tissues (controls from healthy patients) were obtained with informed consent and ethics persmission from patients who had surgery at the Department of Visceral, Thoracic and Vascular Surgery at the University Hospital Carl Gustav Carus of TU Dresden. Organoids were established following a previously published protocol ^32^. WNT2 gene amplification was determined by whole genome sequencing with HighSegXten (Illumina).

### Growth factor withdrawal experiments in gastric corpus organoids

All organoids were maintained in complete medium and passaged at day 0 at a 1:5 ratio. Individual wells were supplied with either complete medium or selection medium deficient in RSPO1 or WNT3A supplement. Medium changed every other day and organoids were passaged every week until the end of the experiment.

For drug treatments, all organoids were maintained in complete medium before treatment. At day 0, organoids were passaged and maintained in selection medium containing 10mM Wnt-C59 (Selleckchem), 50μM SR-11302 (Tocris), or 50μM T-5224 (Tocris) until the end of the experiment. Medium was refreshed every two days and organoids were passaged every week.

#### HER2 3+ gastric cancer organoid growth factor withdrawal experiment

6 HER2 3+ gastric cancer organoids were cultured in Wnt-free normal culture medium supplemented with C59 and/or 100nM Wnt. The culture medium was changed every 3 days. Organoids were passaged at a ratio of 1:3 every 8-9 days, when the Matrigel dome containing the organoids reached ∼90% confluency. After passage, 10uM ROCK inhibitor (Tocris) was added.

### CRISPR-mediated Tp53 targeting in gastric corpus organoid

Gastric corpus organoids were established from Anxa10-CreERT2-Cdh1^f/f^-Kras^G12D^ mice and the *Tp53* allele was targeted by co-transfection with sgRNA (GTGTAATAGCTCCTGCATGGGGG) together with the Cas9 expressing vector using a NEPA21 electroporator. Successfully targeted organoids were selected with 1 mM Nutlin-3 (Selleck Chem) treatment for 2 weeks with passaging every week.

### Targeting of hKRAS^G12D^ in human gastric organoid

KRAS alleles were targeted using two plasmids. For this purpose, two sgRNA sequences for KRAS as well as an sgRNA sequence to linearize the repair template plasmid were inserted into the px458_Conc3 plasmid (Addgene Plasmid #134450) using Golden Gate Cloning, as described by Andersson-Rolf et al., 2016. The generated plasmid, together with a repair template plasmid introducing the G12S point mutation, was transfected by electroporation following the protocol of Gaebler et al., 2020. In addition to the G12S point mutation, the repair template also contained several silent mutations designed to prevent the sgRNAs from binding again. Successfully transfected organoids were selected by EGF deprivation for four weeks, with passaging every week. The integration of the repair template at the correct position in the KRAS gene was confirmed by PCR with the Q5 polymerase. The PCR primers were designed so that one primer binds outside the repair template and the other binds over the silent mutations of the repair template. Primer sequences are listed below.

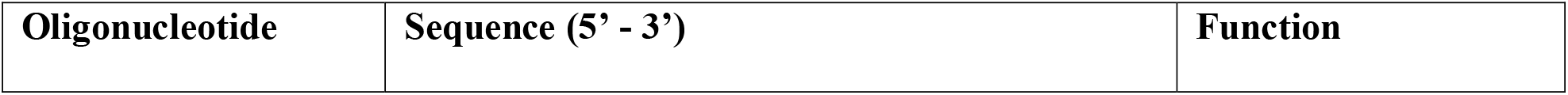
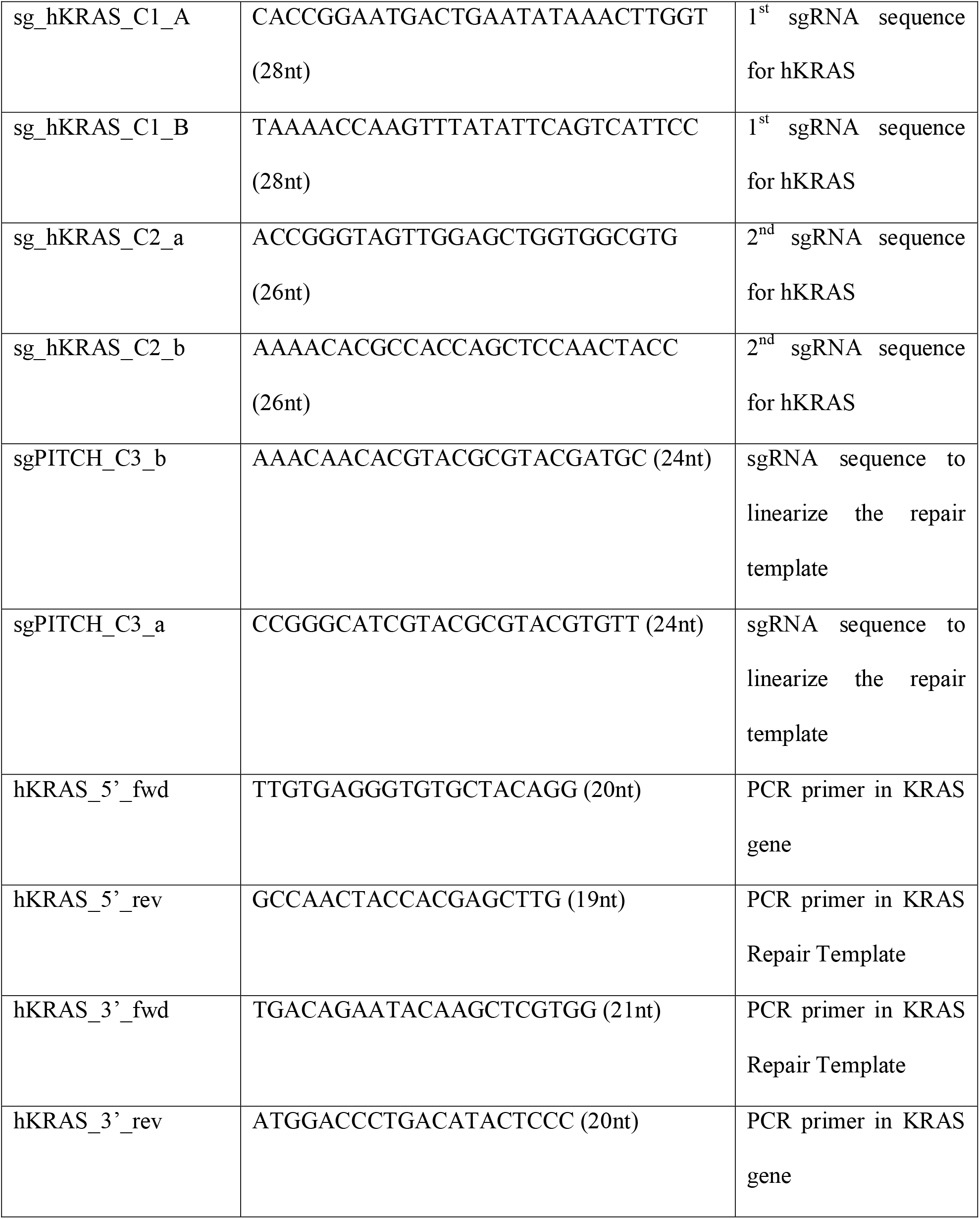

### Single-cell multi omics and gene expression sample preparation

Mouse gastric corpus organoids were maintained in complete medium (wild-type) or RSPO1 deficient medium (Ax10-RZ and Ax10-RZK) until 10 days before the experiment. The organoid culture medium was replaced to complete medium for all samples and maintained for a week after which organoids were passaged 3 days before the experiment. Single-cell dissociation was carried out by incubating mechanically disrupted organoids with TrypLE for 30 min at 37 °C. Nuclei from single cells were extracted. Library preparation and sequencing were performed by Vienna BioCenter Core Facilities. The libraries from each sample were sequenced using an Illumina NovaSeq system.

### Single-cell multiome (RNA-seq and ATAC-seq) data processing

To generate count matrices consist of both gene expression and chromatin peaks for each organoid sample such as Rnf43/Znrf3 double knock-out (RZ), and Kras activated RZ-DKO (RZK), cellranger-ARC count (v6.1.1)^37^ was utilized with default option and mouse reference (‘refdata-cellranger-arc-mm10-2020-A-2.0.0’) provided in 10x Genomics. Based on both Signac (v1.8.0)^38^ and Seurat (v4.1.0) ^39^ pipelines, we generated analytic objects from ‘filtered_feature_bc_matrix’ and ‘atac_fragments.tsv.gz’ of cellranger based on mouse genome which is output from ‘GetGRangesFromEnsDb’ with ‘EnsDb.Mmusculus.79’ for annotations. We filtered out poor-quality cell nuclei with unique molecular identifiers (UMIs) for RNA less than 1,000 and more than 25,000, and for ATAC less than 1,000 and more than 100,000, percent mitochondrial genes more than 15%, and nucleosome signal more than 3 for both samples. Using the selected nulei, we sequentially conducted NormalizeData (log-normalization) and FindVariableFeatures (‘vst’ method, 2,000 features) functions for RNA expression profiles. We also implemented FindTopFeatures with min.cutoff is ‘q0’, RunTFIDF, and RunSVD for both ATAC and macs2 assays for peak profiles from Signac package. After preprocessing individual objects from RZ-DKO and Kras activated RZ-DKO samples, we performed SCTransform for RNA expression profile to prepare integration for the objects. The listed objects were integrated with the sequential functions such as ‘SelectIntegrateionFeatures’, ‘PrepSCTIntegration’, ‘FindIntegrationAnchors’, and ‘IntegrateData’. To generate cluster map for the integrated object, we performed ‘ScaleData’, ‘RunPCA’, ‘RunUMAP’, ‘FindNeighbors’, and FindClusters (‘resolution’ is 0.75, Louvain algorithm) with 30 PCs.

### Single-cell multiome (RNA-seq and ATAC-seq) data analysis

In order to annotate cell types in the dataset, we used cell type markers such as Muc6, Gkn3 for neck cells, Lgr5 for Lgr5 positive cells, Tff2, Glipr1, CD44, Cftr for SPEM, Wnt7a, Wnt7b for Wnt7 positive cells, Mki67, Stmn1 for proliferating cells, and Muc5ac, Tff1, Gkn1, Gkn2 for pit cells. We utilized ‘DotPlot’ from Seurat package for Fig. 3B and ‘dittoBarPlot’ function in dittoSeq R package (v1.6.0)^40^ for Fig. 3C with corresponding cell type markers and cell type annotation. We defined the Wnt7b expressing cells for Fig. 3E which have non-zero unique molecular identifier (UMI) count of RNA count-matrices for Wnt7b. For identification of differentially expressing genes, ‘FindMarkers’ function was utilized. Based on JASPAR2020 dataset^41^ and macs2^42^ peaks, we identified differentially accessible peaks (DAPs) and enriched motifs for each DAP with adjacent p-value < 0.005.

## Acknowledgements

We would like to thank Anne Gocht for pilot experiments for this project.

## Supplementary information

**Supplementary Table 1**. PCR primer sequences for quantitative real-time PCR and genotyping PCR, Related to Figure 1 and Extended Data 2

Correspondence and requests for materials should be addressed to Daniel E. Stange, and Bon-Kyoung Koo.

---

**Extended Data 1.**
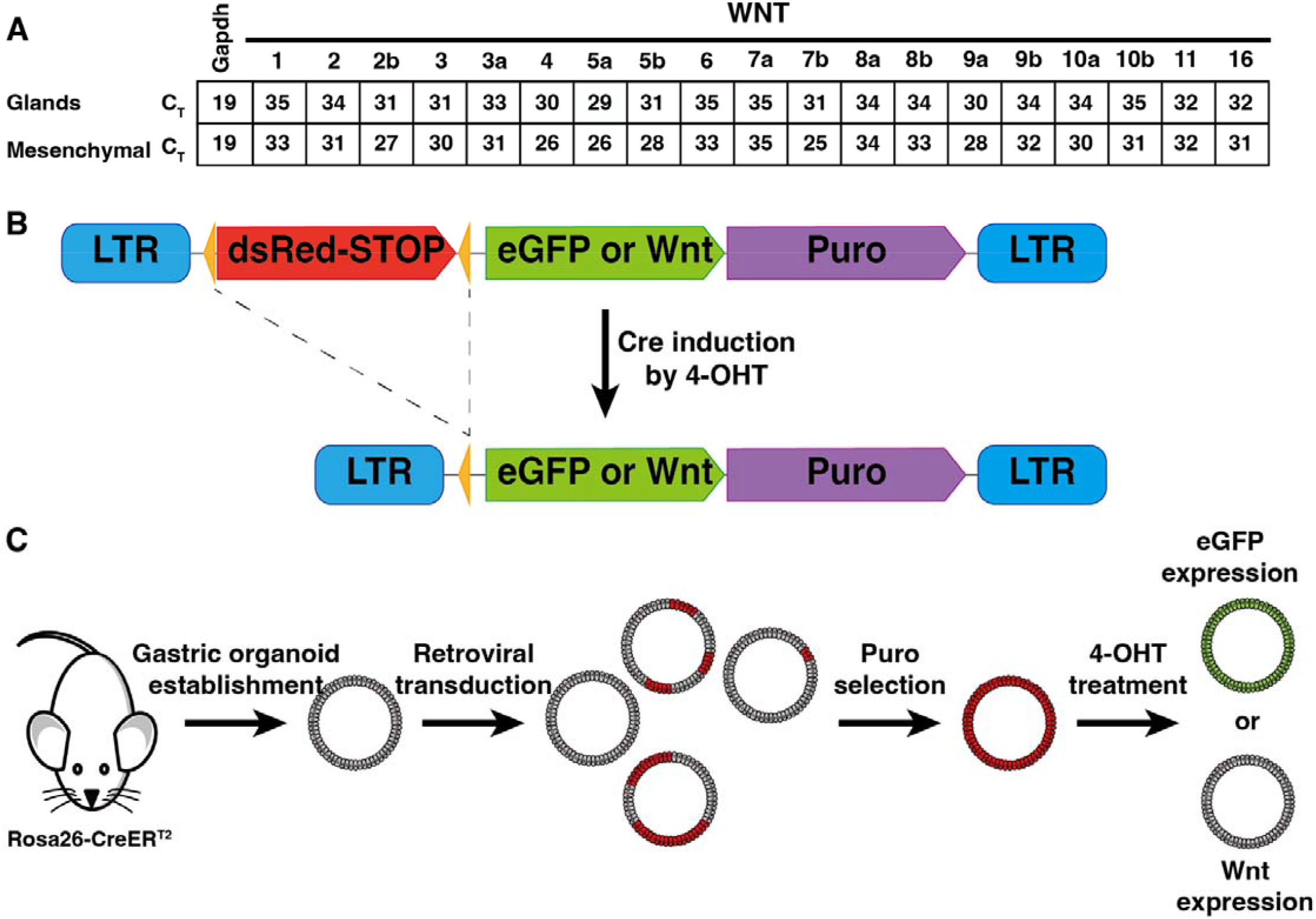
Retrovirus-mediated gene overexpression in gastric corpus organoids. (A) Calculated C_T_ values for expression of a panel of WNT genes in gastric glands and mesenchymal compartment. (B) Schematic representation of Cre-mediated recombination for the eGFR or WNT gene expression (C) Diagram of experimental workflow used to generate retrovirus-mediated WNT gene overexpression in gastric corpus organoids

**Extended Data 2.**
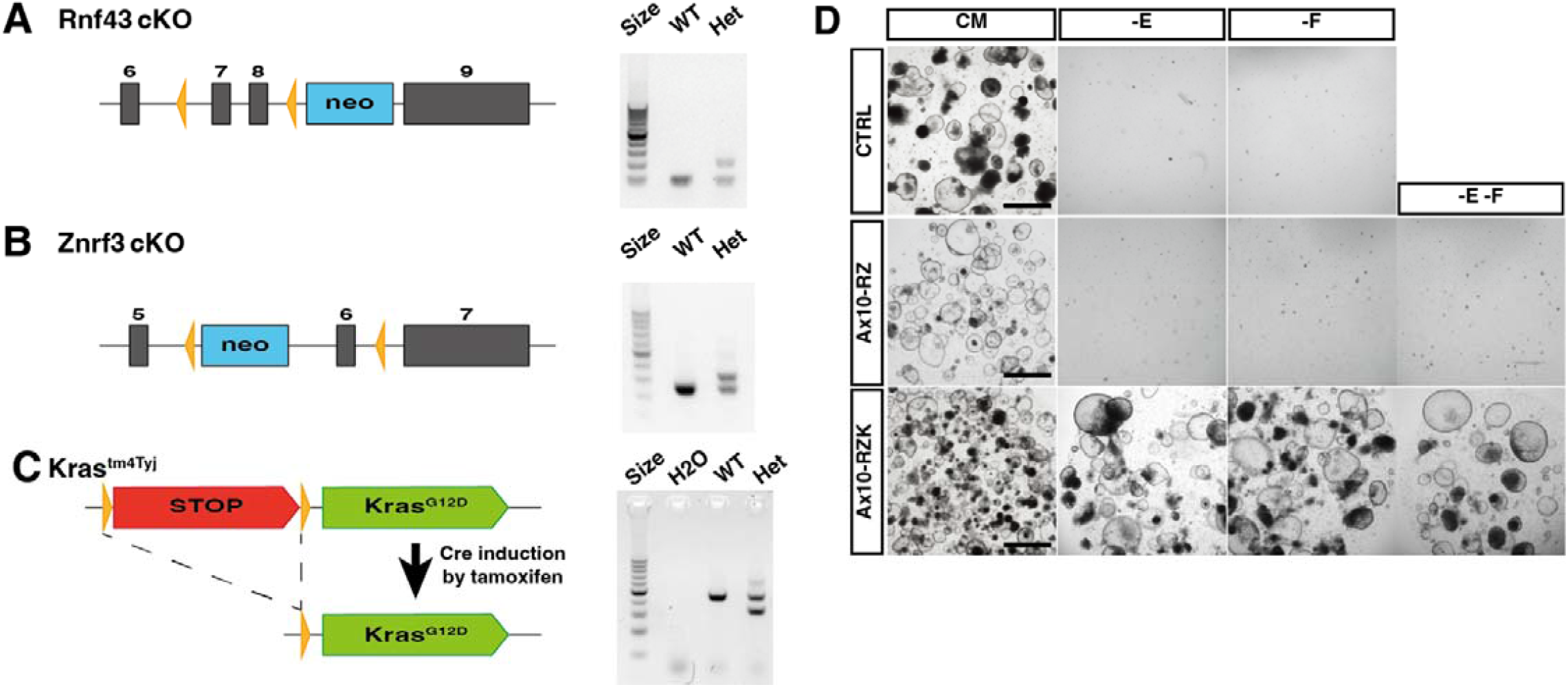
Generation and genotyping of mutant mouse lines. (A) Schematic representation of Rnf43^f/f^ allele generation. Right: genotyping gel. (B) Schematic representation of Znrf3^f/f^ allele generation. Right: genotyping gel. (C) Schematic representation of lsl-Kras^G12D^ (Kras^Tm4Tyj^) allele generation. Right: genotyping gel. (D) Niche requirements for CTRL, Ax10-RZ and Ax10-RZK gastric organoids. Organoid growth was examined after 6 passages. Healthy organoid growth is cystic with a clear center. Shown are representative images of organoids isolated from 2-4 mice per genotype. CM, complete medium; -E, WNRF; -F, WENR; -E-F, WNR medium. Scale bars, 1000 μm

**Extended Data 3.**
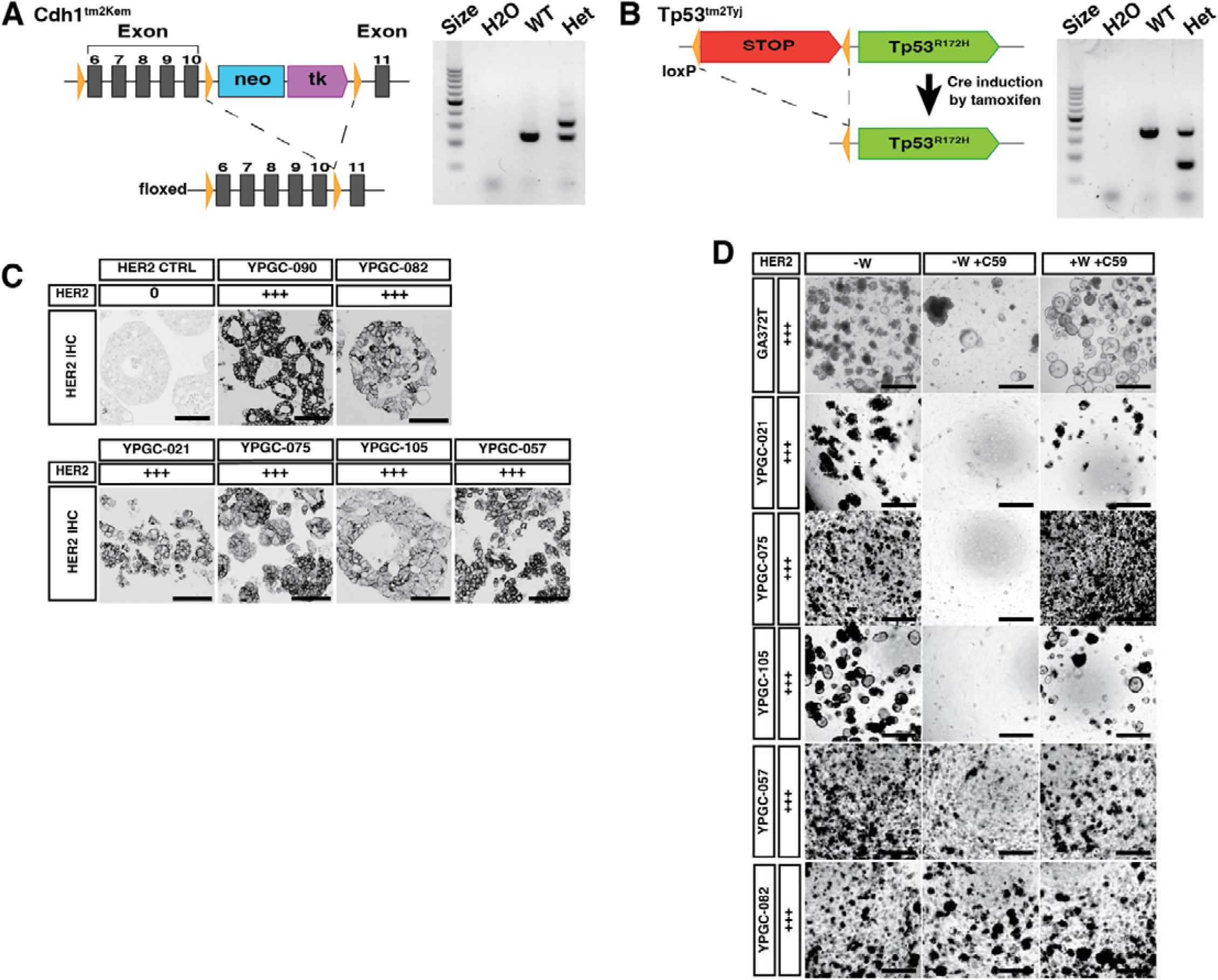
Alternative routes to WNT independence in mouse and relevance to human gastric cancer patients. (A) Schematic representation of Cdh1^f/f^ (Cdh1^tm2Kem^) allele generation. Right: genotyping gel. (B) Schematic representation of Tp53^f/f^ (Tp53^Tm2Tyj^) allele generation. Right: genotyping gel. (C) HER2 immunohistochemistry of corpus organoids derived from a human gastric cancer patient. HER2 levels are classified as negative (0), low-grade (+) and positive (+++) based on guidelines published by Bartley et al., 2016. Scale bars, 100 μm. (D) Niche requirements for human gastric cancer patient-derived organoids. Organoid growth was examined after 6 passages. Healthy organoid growth is cystic with a clear center or growing in grape-like structures. CM, complete medium; -W, ENFR; -W+C59, ENFR medium with C59 (10mM); +W+C59, WENFR with C59 (10 mM). Scale bars, 1000 μm.

